# A new means of energy supply driven by terahertz photons recovers related neural activity

**DOI:** 10.1101/2022.11.22.517481

**Authors:** Xiaoxuan Tan

## Abstract

Continuous and efficient energy capture represents a long-sought dream of mankind. The brain is a major energy consuming organ, an adult brain accounts for about 2% of the body weight but consumes about 20% of the body’s energy. However, it is still unclear how the brain achieve efficient use of energy. Here, using nerve cells as test subjects, we found that THz photons with a specific frequency can effectively restore the reduced frequency of action potentials caused by inadequate adenosine 5’-triphosphate (ATP) supply, which has been demonstrated as a novel mode of energy supply, present photons emission at a particular frequency from the breaking of the ATP phosphate bond. This energy supply mechanism may play a key biophysical basis for explaining how the body efficiently obtain energy, because the quantized chemical reactions could have a high energy efficiency and ultrahigh selectivity compared with the traditional thermochemistry and photochemistry.

## Introduction

As a universal energy-storing molecule, adenosine 5’-triphosphate (ATP) is used by virtually all living organisms, and plays an important role in both physiological and pathophysiological processes[1-3]. In addition, ATP also regulates many brain functions such as synaptic transmission[4], glutamate release[5], pain sensation[6] and have complex interactions with other signaling such as reactive oxygen species (ROS) or calcium[7-9]. Despite ATP is important for both health and disease, the detailed mechanisms underlying the efficient energy supply of ATP are poorly understood, especially *in vivo*.

It has become increasingly popular for people to take some foods or drugs as functional beverage that are expected to obtain energy to achieve the purpose of maintaining the efficient work of the human body for a long time. However, functional energy foods have very limited energy supply, and stimulants are banned in most cases because of their side effects[10,11]. Early observations have proposed that human high intelligence is involved in spectral redshift of biophotonic activities in the brain because biophoton spectral redshift could be a more economical and effective measure of biophotonic signal communications and information processing in the brain[12,13], The improvement in cognitive function is driven by the need for more effective communication that requires less energy, but, so far, there is no direct experimental evidence for the role of photons in brain information processing and energy supply.

A recent study proposed the concept for the quantized chemical reaction resonantly driven by multiple FIR/MIR photons[14-16]. Theoretic studies revealed that THz photons released by ATP phosphate bond breakage could drive life activities. Nevertheless, the quantum mechanisms underlying the THz bioeffects remain elusive, and the experimental evidences are still lacking. In this work, we employed nerve cells as test object to study the effects of THz photons on energy supply. We surprisingly found that without introducing any exogeneous substances and implanting electrodes, THz photons could effectively recover reduced action potential emission frequency due to inadequate ATP supply, and we ruled out the possibility of restoring ATP production for energy supply (Fig.1A). The phenomenon can clearly demonstrate that THz photons can regulate brain information processing and energy metabolism directly.

**Fig.1.**
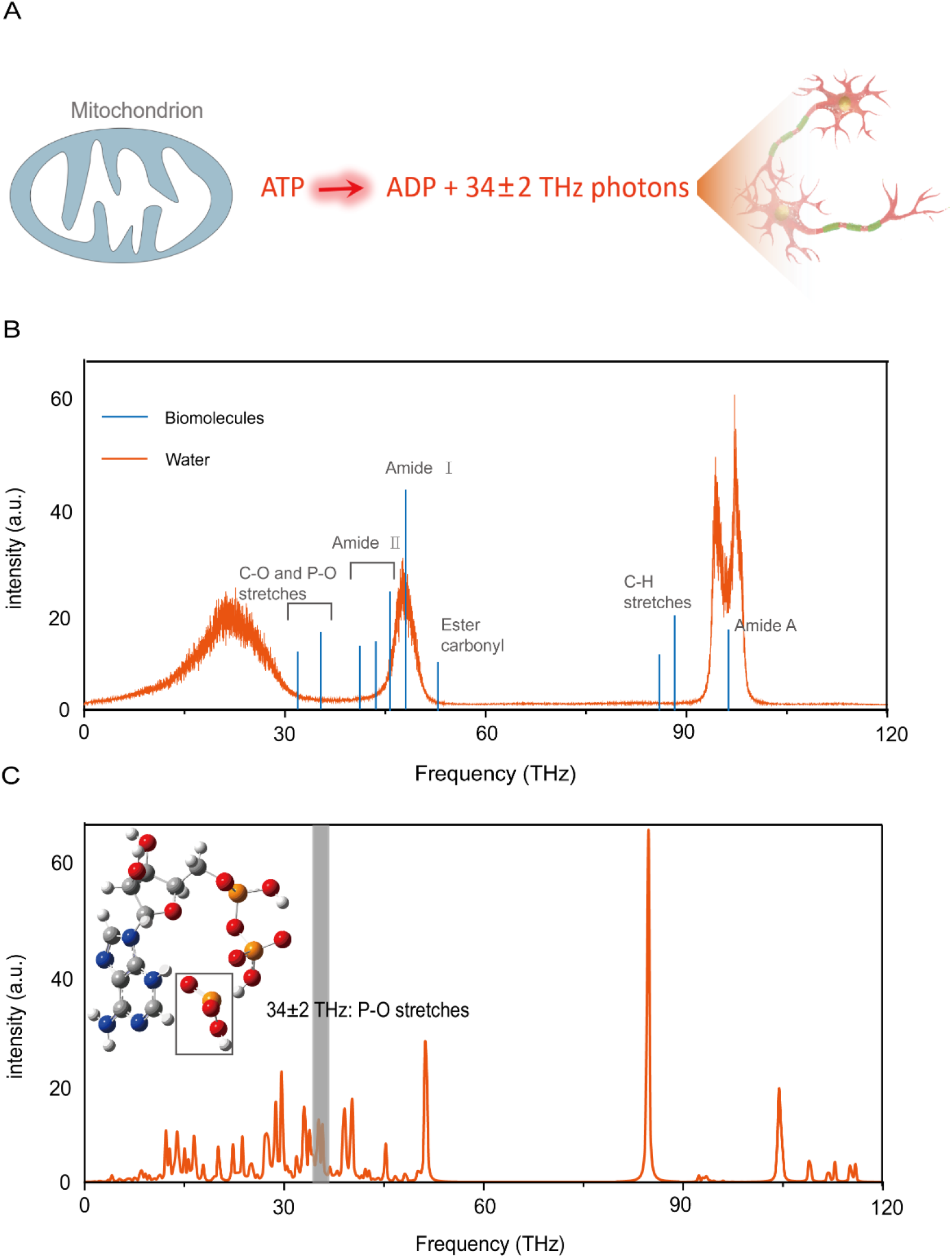
Mechanistic study of the energy supply driven by THz photons. (**A**) Schematic indicating the process of THz photons release and it acted on nerve cells. (**B**) Theoretic absorption spectra of the bulk water model (orange curve) and the THz photons absorption spectra of biomolecules (blue curve). (**C**) Theoretic absorption spectra of the ATP. Inset, schematic illustration of the P-O stretches around 34±2 THz.

## Results and discussion

Photons with a specific frequency can participate in multiple life activities such as ion channel dynamics and DNA unwinding through changing vibration levels of a confined molecule excited (Fig.1B)[17-20]. Previous studies have proposed that the principle of terahertz wave energy supply is the release of 34 ± 2 THz photons by phosphate bond break, then photons acts as a massless reagent to supply energy[15,16]. We therefore calculated whether the IR vibration spectra of ATP have strong formants at 34 ± 2 THz by density functional theory (DFT). As Fig.1C shown, the ATP shows significant THz absorption in this range. In particular, ATP supplies energy by breaking high-energy phosphate bonds and the THz frequency (34 ± 2 THz) indeed could resonate with the vibration of phosphate bond. These data agree with the previous results and suggest that 34±2 THz may well be the direct energy source for life activities such as biosynthesis.

Next, we designed and fabricated terahertz lasers that can be used to irradiate biological tissues. The λ ∼ 8.6 μm quantum cascade laser (QCL) wafer was grown on an n-doped (Si, 2 × 10^17^ cm^−3^) InP substrate by solid source molecular beam epitaxy (MBE) based on a slightly-diagonal bound-to-continuum design, as is identical to that in Ref. 21 (for detailed protocol, see Supplemental Methods). As Fig.2 shown, the power of QCL has good portability and easy integration. The laser output by QCL is coupled into the fiber by the coupler for collimation. The fiber (PIR600/700-100-FC/PC-FT-SP30) with a core diameter of 200 μm and a numerical aperture (NA) of 0.35±0.05 we used is flexible, so there is good flexibility to change the stimulation target. We delivered the THz laser with the following parameters: average irradiation power 30 ± 2 mW (measurements were made before and after each experiment to ensure stable output), pulse width 2 us, and repetition rate 200 kHz (for detailed protocol, see Supplemental Methods).

**Fig.2.**
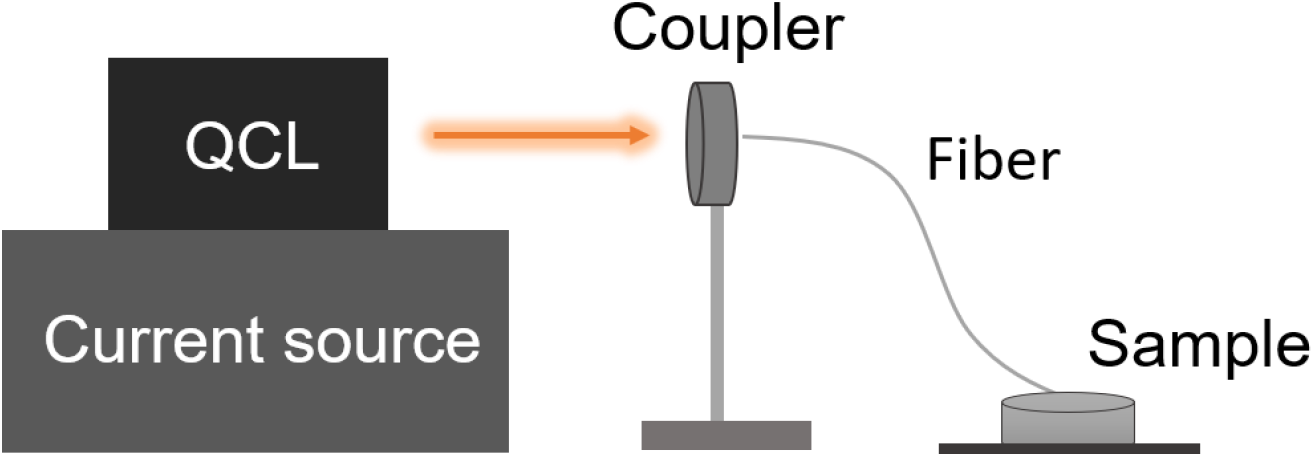
Schematic drawing of the THz illumination device. The QCL was described in detail in a previous report^[21]^. The biological tissue such as brain slice is incubated in a chamber containing perfusion solution (also see SI methods).

To determine whether 34 ± 2 THz photons have the function of energy supply, we first asked whether 34 ± 2 THz photons could restore the diminished life activities caused by insufficient energy supply. To answer this question, we took neurons as the test object (layer-5 pyramidal cells in acute slices of mouse prefrontal cortex) and performed patch-clamp experiment to quantify the firing frequency of neurons’ action potentials. We first inhibited mitochondrial ATP generation and electron transfer pathways by using mitochondrial ATP production inhibitor (mito-I) composed of rotenone (mitochondrial complex I blocker, applied to inhibit cyclophilin D to reduce mPTP activity in cells, 10 μM), anti-mycin A (to inhibit electron transfer between Cyt b and Cyt c of complexIII, 4 μM) and oligomycin (to inhibit ADP phosphorylation and electron transport, 10 μM)[22]. The experimental results show that mito-I we selected can effectively block the energy supply of neurons. As Fig.3A shown, about 2 minutes after the addition of mito-I to artificial cerebrospinal fluid (ACSF), the neurons were injected with 400 pA currents and the firing frequency of action potential (AP) was significantly lower than that before the addition of mito-I (from 20.67 ± 1.33 to 13.67 ± 2.09 Hz, *n* = 6, *t*_*5*_ = 4.134, *P* = 0.0090, paired Student’s *t* test; Fig. 3B). When THz applied, the AP frequency increased significantly from 13.67 ± 2.09 to 17.00 ± 1.84 Hz (*n* = 6, *t*_*5*_ = 5.000, *P* = 0.0041, paired Student’s *t* test; Fig. 3B), basically returned to the level before adding mito-I (from 20.67 ± 1.33 to 17.00 ± 1.84 Hz, *n* = 6, *t*_*5*_ = 2.314, *P* = 0.0686, paired Student’s *t* test; Fig. 3B). Although slightly reduced, mito-I had no significant effect on the AP amplitude (from 80.92 ± 3.54 to 74.94 ± 6.33 mV, *n* = 6, *t*_*5*_ = 2.089, *P* = 0.0910, paired Student’s *t* test; Fig. 3B) and halfwidth (from 1.71 ± 0.17 to 1.61 ± 0.32 ms, *n* = 6, *t*_*5*_ = 0.6535, *P* = 0.5423, paired Student’s *t* test; Fig. 3B). Meanwhile, although both mito-I and THz illumination led to a slight decrease in the AP area (from 142.4 ± 5.65 to 118.0 ± 8.45 mV·ms, *n* = 6, *t*_*5*_ = 3.937, *P* = 0.0110 and from 118.0 ± 8.45 to 78.03 ± 13.79 mV·ms, *n* = 6, *t*_*5*_ = 2.716, *P* = 0.0420, respectively, paired Student’s *t* test; Fig. 3B), their kinetic properties such as rise slope and decay slope did not change significantly (Fig. S1).

**Fig.3.**
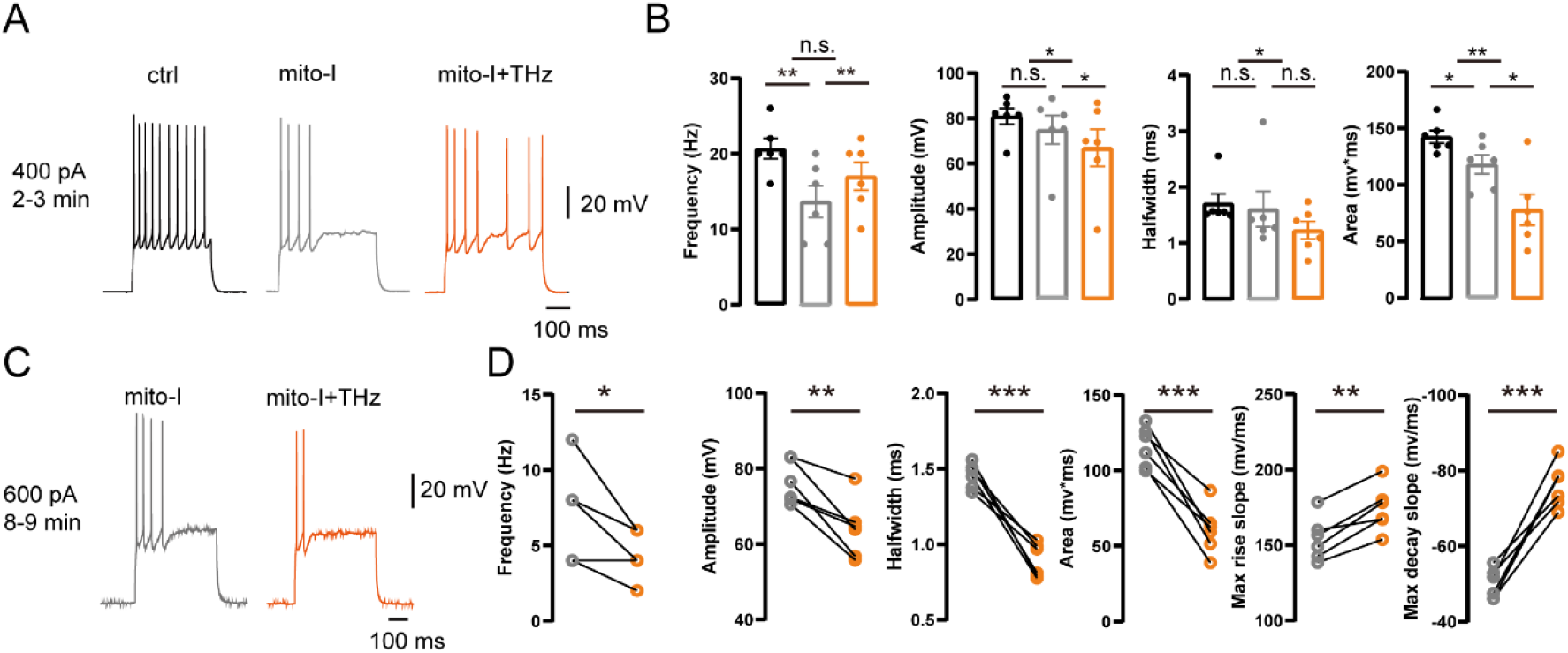
terahertz photons recovers related neural activity. (**A**) Representative AP spiking before adding mito-I, after adding mito-I and THz illumination after adding mito-I (2-3 min). (**B**) Comparison of the changes in frequency, amplitude, halfwidth and area, the three groups of data of each figure correspond to the three states of the Fig.3A. (**C**) Representative AP spiking before and during THz illumination (8-9 min). (**D**) Comparison of the changes in frequency, amplitude, halfwidth and area, the two groups of data of each figure correspond to the two states of the Fig.3C. *, **, *** and **** represent P < 0.05, 0.01, 0.001 and 0.0001, respectively. Repeated measurement ANOVA. Error bars represent s.e.m.

The effect of mito-I accumulated over time, about 8 minutes after the addition of mito-I to ACSF, the neurons no longer generated APs when 400 pA currents were injected. Therefore, we injected a much stronger 600 pA current (Fig. 3C). Interestingly, at this point the frequency of AP decreased from 7.33 ± 1.23 to 4.67 ± 0.67 Hz during THz illumination (Fig. 3D), the AP amplitude and halfwidth were also dramatically reduced from 76.26 ± 2.30 to 64.04 ± 3.17 mV during THz illumination (Fig. 3D) and from 1.46 ± 0.03 to 0.89 ± 0.05 ms during THz illumination (Fig. 3D), which resulted in the area decreased from 115.9 ± 5.49 to 60.74 ± 6.48 mV·ms (Fig. 3D). Meanwhile, dynamic properties such as rise slope and decay slope were significantly enhanced (from 154.2 ± 5.89 to 174.3 ± 6.26 mV/ms, *n* = 6; from -50.01 ± 1.64 to -76.04 ± 2.39 mV/ms, respectively; Fig. 3D). We speculate that this is because the AP is affected by ATP and other factors, such as Na^+^ - K^+^ pump activity, ion concentration inside and outside the membrane, enzyme activity, membrane properties such as membrane resistance, etc[23-26]. Long periods of inadequate energy supply leads to irreversible changes or damage to a variety of influencing factors (Fig. S3), under the complex and comprehensive action of many factors, THz illumination at this time enhanced the dynamic properties of AP but did not restore the AP frequency. Besides, Our previous study found that THz radiation can change the kinetic properties of ion channels and different cell states lead to different excitability changes[19,20], so we speculate that THz may enhance the kinetic properties of Na+ channels in this cellular state, leading to an increase in the rising slope of the AP, which also accelerate the repolarization process, leading to the decrease in the AP amplitude.

Does THz illumination change the amount of ATP in nerve tissue? We next employed two-photon Ca^2+^ imaging to visualize the changes of ATP during THz illumination *in vivo*. We measured ATP in neurons and astrocytes respectively in mouse sensory cortex. The sensory cortex was injected with an ATP-sensitive probe (rAAV-hSyn-GRAB (ATP1.0), target neurons or rAAV-GfaABC1D-GRAB (ATP1.0), target astrocytes). We performed real-time (40 Hz) two-photon imaging over the entire course of THz illumination (2 mins before, 4 mins THz illumination, 4 mins after; Fig. 4A). Since the two fluorescence probes we used have just been developed and no relevant reports have been reported[27], we first tested the effectiveness of the two probes. As the Fig. S1 shown, after the addition of mito-I (the drug composition was the same as that of patch clamp experiment, and the concentration was 5 times that of brain slice experiment), the fluorescence intensity decreased significantly and after the addition of ATP (100 μM), the fluorescence intensity increased, which support the effectiveness of the fluorescent probes we used.

**Fig.4.**
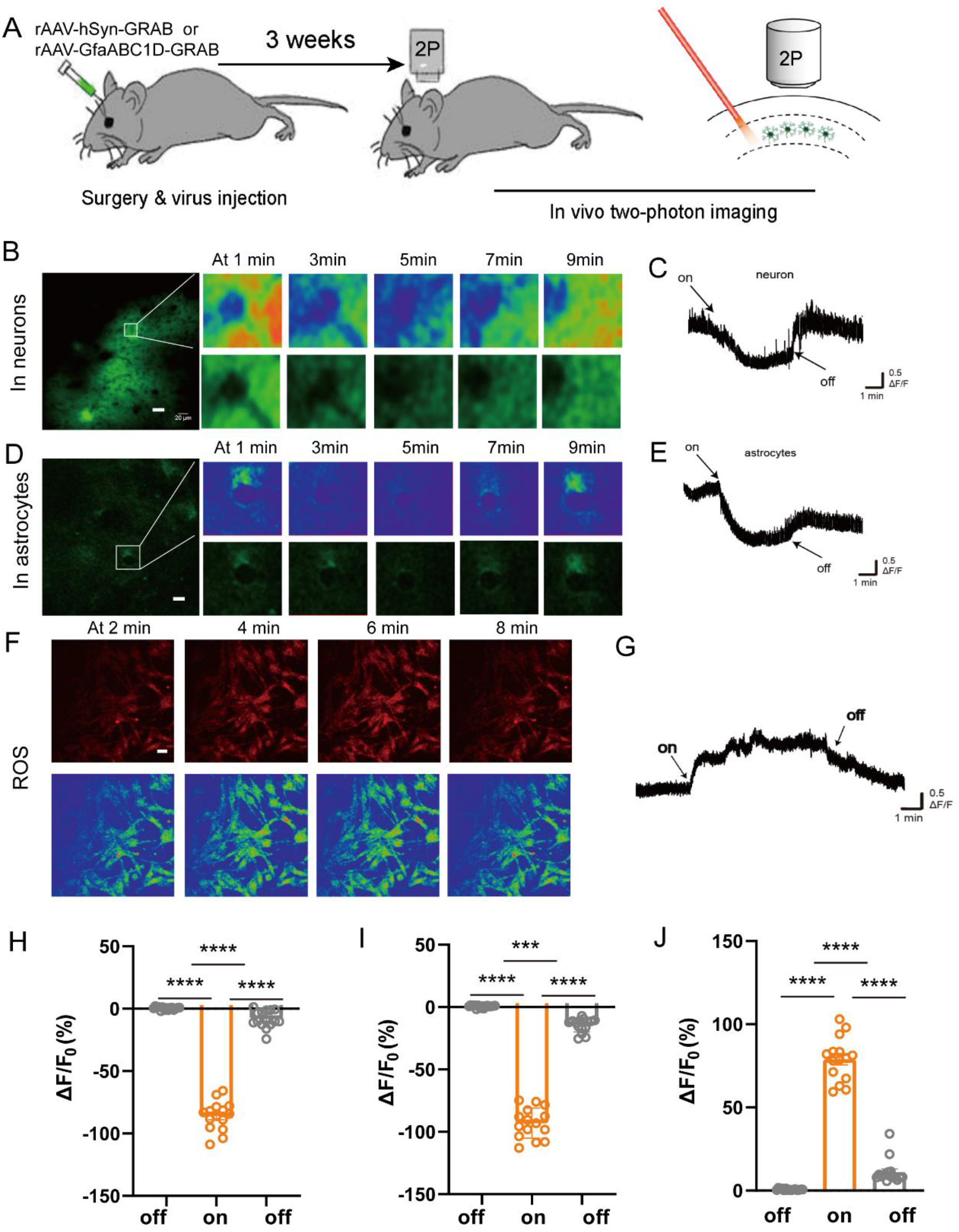
THz illumination leads to a decrease in ATP and an increase in ROS. (A) Schematic diagram depicting the experimental protocol in which an AAV encoding promoter is injected into the mouse sensor cortex, followed by two-photon imaging. (B and C) Exemplar fluorescence images, pseudocolor images, and individual traces of the fluorescence response of ATP measured in neurons (THz illumination at 4-6 minutes). (D and E) Exemplar fluorescence images, pseudocolor images, and individual traces of the fluorescence response of ATP measured in astrocytes (THz illumination at 4-6 minutes). (H) The summary data of ATP in neurons represent 15 ROIs from 5 mice. (I) The summary data of ATP in astrocytes represent 15 ROIs from 5 mice. (J) The summary data of ROS represent 15 ROIs from 3 coverslips. Scale bars represent 20 μm. *, **, *** and **** represent P < 0.05, 0.01, 0.001 and 0.0001, respectively. Repeated measurement ANOVA. Error bars represent s.e.m.

Next, we used the above two probes to test the direct effects of THz illumination on ATP in neurons and astrocytes *in vivo* via two-photon imaging. An example neuron (Fig. 4B, C) showed that THz illumination reliably induced ATP in neurons during the illumination time (2-6 min) and astrocytes changed in the same way as neurons (Fig. 4D, E,H,I). The changes of intracellular ATP in HEK293T cells is consistent with the extracellular ATP in vivo (Fig. S3)[28,29]. We speculate that the reason is that the body may have a “compensatory” response, that is, the THz photons are energy-supplying during the illumination, so the cells no longer need to synthesize large amounts of ATP, resulting in reduced ATP release. We also

Previous studies have reported that ATP could play the role of peroxidase in mitochondria to remove excess reactive oxygen species (ROS) and ruled out the possibility that ATP promoted the decomposition of ROS by releasing energy[30]. We therefore tested whether and how THz illumination in the nerve cells affect the ROS in mitochondria. We used a mito-targeted reporter (MitoSOX Red, 5 μM) to measure mitochondrial superoxide production, imaging time was 10 min for each group (2 mins before, 4 mins THz illumination, 4 mins after)[31]. A typical example of changes of ROS in mitochondria is shown in Fig. 4F&G, in which there is obvious increase of ROS during THz illumination (Fig. 4J). These results together suggest that THz illumination results in a decrease of ATP in nerve cells and affects the biochemical reactions involved in ATP such as the breakdown of ROS.

## Conclusion

In this study, we report that THz photons can effectively recover reduced action potential emission frequency due to inadequate ATP supply and change the amount of ATP and ROS in nerve cells without introducing any exogeneous substances. Besides, this control technology has good targeting and direction of brain tissue *in vivo*, which is fundamentally different from some well-known and widely-deployed human function regulation technology such as optogenetics and electrode implantation. Our results illustrate that biophotons may play a key role in neural information processing and encoding and may be involved in quantum brain mechanism.

We hope that our findings can bring a new viewpoint that neural communication or physiological signal transduction is likely to be essentially the transmission of endogenous terahertz waves and other high-frequency electromagnetic signals and provide an explanation of how energy can be used efficiently. However, due to the complex coherence of biological quantum information[32-36], its related exploration still has a long way to go.

Overall, if the generation, transmission, and coupling of neural signals are in the form of THz photons, then exogenous THz photons input can change and even regulate various life activities, which can not only explain the efficient and low-consumption information processing of human beings, but also provide new ideas for exploring the mysteries of the brain and the development of artificial intelligence. At the same time, the relevant technical means have great application prospects in the fields of disease treatment and human function regulation.

## Acknowledgements

This work was supported by the National Natural Science Foundation of China (No.12225511 and T2241002), National Defense Science and Technology Innovation Special Zone, and the National Supercomputer Center in Tianjin. We thank the support from Prof. Yulong Li and Dr Zhaofa Wu of Peking University. C.C. acknowledges the support from the XPLORER Prize. The authors are grateful to Dr. Bo Song for inspiring our work.

## Competing interests

The authors declare no competing interests.

## Supporting information

## Methods

### Animals

C57BL/6J male mice (2-3 months old) were provided by the Laboratory Animal Center of the Qsinghua University. All experimental procedures were performed in accordance with institutional animal welfare guidelines with the approval of Qsinghua University Animal Care and Use Committee.

### THz-QCL laser

We used a quantum cascade terahertz laser for this study, as describe in previous studies[1,2]. The quantum cascade laser (QCL) wafer was grown on an n-doped (Si, 2 × 10^17^ cm^−3^) InP substrate by solid source molecular beam epitaxy (MBE) based on a slightly-diagonal bound-to-continuum design. The epi-wafer was etched into double channel geometry with a ridge width of 9 μm. After etching, Fe-doped semi-insulating layers were regrown by metal-organic chemical vapor deposition (MOCVD). Silicon oxide was grown by plasma-enhanced chemical vapor deposition (PECVD) as electrical isolation layer. We set a 3 μm wide window for electron injection on each ridge by photolithography and wet etching and ohmic contact was provided by a Ti/Au layer. Then we deposited a Ge/Au/Ni/Au metal contact layer on the backside of the sample after thinning the substrate down to 120 μm. Finally, the sample was high-reflectivity (HR) coated with Al_2_O_3_/Ti/Au/Al_2_O_3_ (200/10/100/120 nm) on the back facets by e-beam evaporation, and then cleaved into 4-mm-long devices. The beam of laser was collimated by an anti-reflectivity (AR) coated high-aperture collimation lens (CL) (LightPath 390037IR1, NA = 0.86, f = 1.87 mm).

### Mouse neucortical slice preparation

Coronal slices of prefrontal cortex were obtained from C57/B6 mice (postnatal ∼2 months). We anesthetized the mice with sodium pentobarbital (10 mg/kg of body weight, intraperitoneal injection) and sacrificed with decapitation. Brain tissues were then immediately dissected out and immersed in ice-cold oxygenated (95% O_2_ and 5% CO_2_) slicing solution in which 126 mM NaCl was substituted by 213 mM sucrose. Slices (250 μm in thickness) were cut with a microtome (VT-1200S, Leica, Germany) and incubated in the aerated normal ACSF containing (in mM): 126 NaCl, 2.5 KCl, 2 MgSO_4_, 2 CaCl_2_, 26 NaHCO_3_, 1.25 NaH_2_PO4, and 25 dextrose (315 mOsm, pH 7.4) and maintained at 34.5 °C for 1.5 h. After incubation, slices were kept in the same solution at room temperature until use.

### Electrophysiological recordings from mouse neocortical slices

Before recordings, the slices were transferred into a recording chamber and perfused with a regular ACSF (0.9 ml/min). Cortical neurons were visualized with an upright infrared differential interference contrast (IR-DIC) microscope (BX51WI, Olympus) equipped with a water-immersed objective (40x, NA 0.8). The patch pipettes for somatic recordings had impedance of 4-6 MΩ. Electrical signals were acquired using a Multiclamp 700B amplifier (Molecular Devices), digitized and sampled by Micro 1401 mk II (Cambridge Electronic Design) at 25 kHz using spike2 software.

### Virus injection

Prior to surgey, the mice were fixed in a stereotactic frame (RWD, Shenzhen, China) under a-combination of xylazine (10mg/kg) anesthesia and ketamine (100mg/kg) analgesia. A volume of ∼150 nl virus was injected using calibrated glass microelectrodes connected to an infusion pump (micro 4, WPL, USA) at a rate of 30 nl/min. The coordinates were defined as dorso-ventral (DV) from the brain surface, anterior-posterior (AP) from bregma, and medio-lateral (ML) from the midline (in mm).

The glass pipettes used to perform virus injection were beveled at 45° with a 10 to 15 μm opening. After infusing solution, we kept the pipette in the brain for about 10 min before pulling up the pipette to avoid backflow during withdrawal. About 150 nl volume of rAAV-hSyn-GRAB (ATP1.0) or rAAV-GfaABC1D-GRAB (ATP1.0) solution (∼2×10^12^ infectious units/ml) was injected into sensory cortex slowly (AP: ∼-1.5 mm, ML: ‘∼2.0mm, DV: ∼0.35mm) for imaging calcium activity with GCaMP6f.

### Two-photon Ca^2+^ imaging

A commercial Nikon A1R two-photon microscope system was used to perform two-photon calcium imaging. Two-photon excitation beam was emitted by a mode-locked Ti: Sa laser (model “Mai-Tai Deep See”, Spectra Physics). We utilized a water-immersion objective (Nikon) with 25 ×/1.10NA to perform imaging. The excitation wavelength was set to 910 nm for Gcamp6f calcium imaging experiments. For somatic imaging, the dimension of field-of-view (FOV) was set 200 μm × 200 μm. We acquired images of 512 × 512 pixels at 30-Hz frame rate. The average power reaching the cortical surface ranged from 30 to 40 mW, depending on the expression efficiency of virus and depth of imaging. Within an imaging time window of ∼3 min (∼1 min per time, three times in total) for each imaging, no sign of photo-damage was observed. Before the experiment, the mice were fixed under microscope objective frequently with the chamber to adapt to the imaging state for real calcium activity.

### Quantum-chemical calculations

Quantum-chemical calculations were performed to obtain the infrared spectroscopy of ATP. The PBE0 exchange correlation functional and 6-311G* basis set were employed in the DFT calculations. The polarized continuum model (PCM) was used for modeling solvation effects together with water as the solvent. The geometry optimization calculations were performed first to obtain the stable conformations of molecules and the lowest energy conformation was used for calculating results. The frequency of absorbing/emissive photon was identified by the vibrational mode. All calculations were performed by the Gaussian 16 software package and Gaussview 6.0 visualization program[3-5].

### Cell cultures

HEK293T cells were obtained from ATCC (CRL-3216) and verified based on their morphology under the microscope and by their growth curve. HEK293T cells were cultured at 37°C in 5% CO2 in DMEM (Biological industries) supplemented with 10% (v/v) fetal bovine serum (FBS, GIBCO) and 1% penicillin-streptomycin (Biological Industries).

Rat primary neurons were prepared from 0-day old (P0) pups (male and female, randomly selected) in Ref.6-7. Cortical and hippocampal neurons were dissociated from the dissected brains in 0.25% Trypsin-EDTA (Gibco) and plated on 12-mm glass coverslips coated with poly-D-lysine (Sigma-Aldrich) in neurobasal medium (Gibco) containing 2% B-27 supplement (Gibco), 1% GlutaMAX (Gibco), and 1% penicillin-streptomycin (Gibco). Based on glial cell density, after approximately 3 days in culture (DIV 3), cytosine β-D-arabinofuranoside (Sigma) was added to the hippocampal cultures in a 50% growth media exchange at a final concentration of 2 uM. The neurons were cultured at 37°C in 5% CO_2_.

### Expression of MaLion in HEK293T cells

For confocal imaging, HEK293T cells were plated on 12-mm glass coverslips in 24-well plates and grown to 60-80% confluence for transfection. Cells were transfected using a mixture containing 1 μg DNA and 1 μg PEI for 4-6 hours and imaged 24-48 hours after transfection.

### Measurement of mitochondrial ROS production

Mitochondrial production of ROS in cells was measured using MitoSOX Red (Invitrogen/Thermo Fisher Scientific), a mitochondrion-specific hydroethidine-derivative fluorescent dye[8]. In brief, cells were incubated with MitoSOX (5 uM; for 30 min, at 37°C, in the dark). The cells were then washed with PBS and MitoSOX fluorescence was measured by confocal imaging.

### Statistical analysis

We used GraphPad Prism 5 and MATLAB to perform the statistical analysis. Except where indicated otherwise, summary data are presented as the mean ± SEM. All measurements were taken from distinct samples.

---

**Fig S1.**
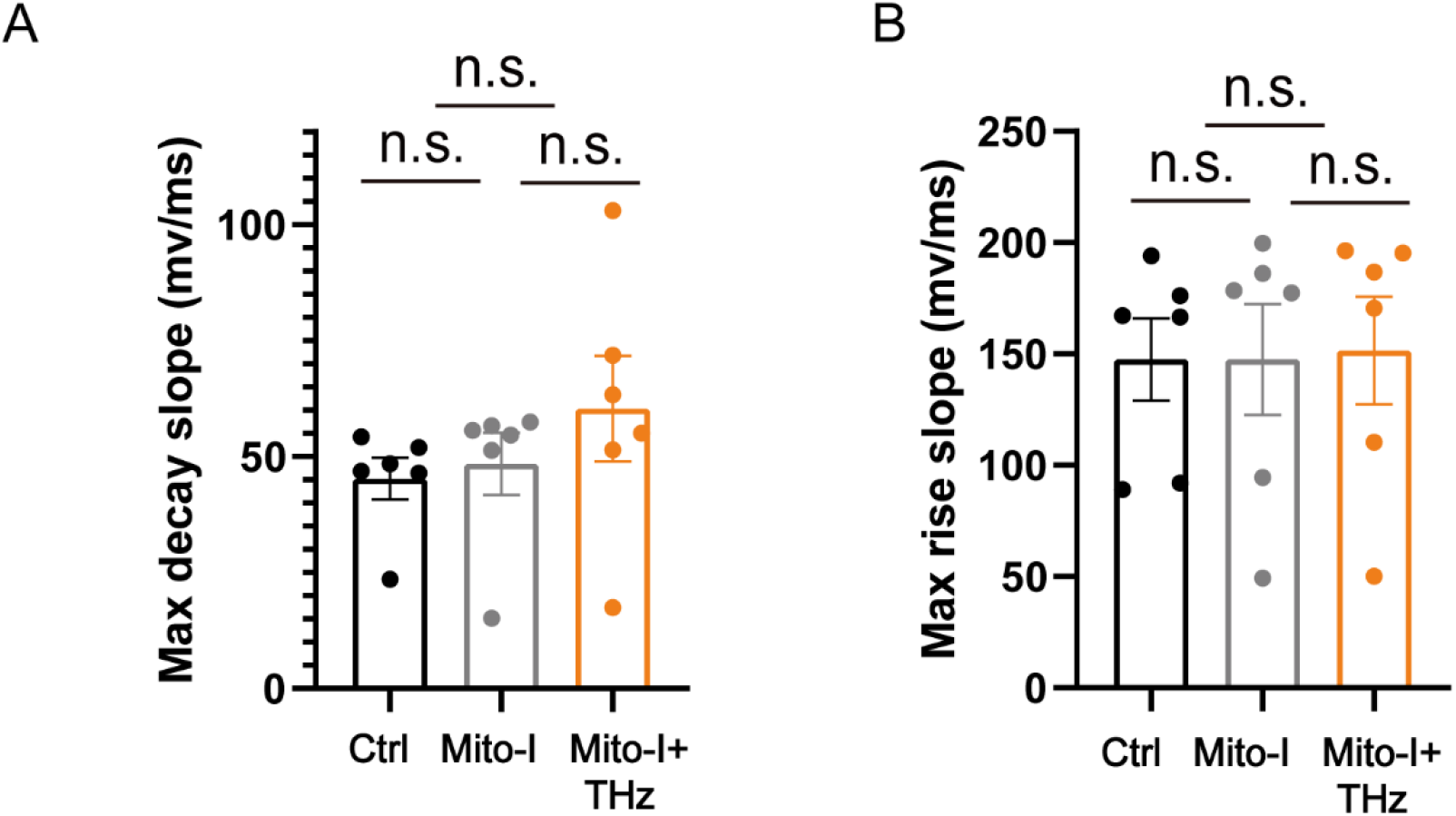
The decay slope (A) and rise slope (B) of the action potential.

**Fig S2.**
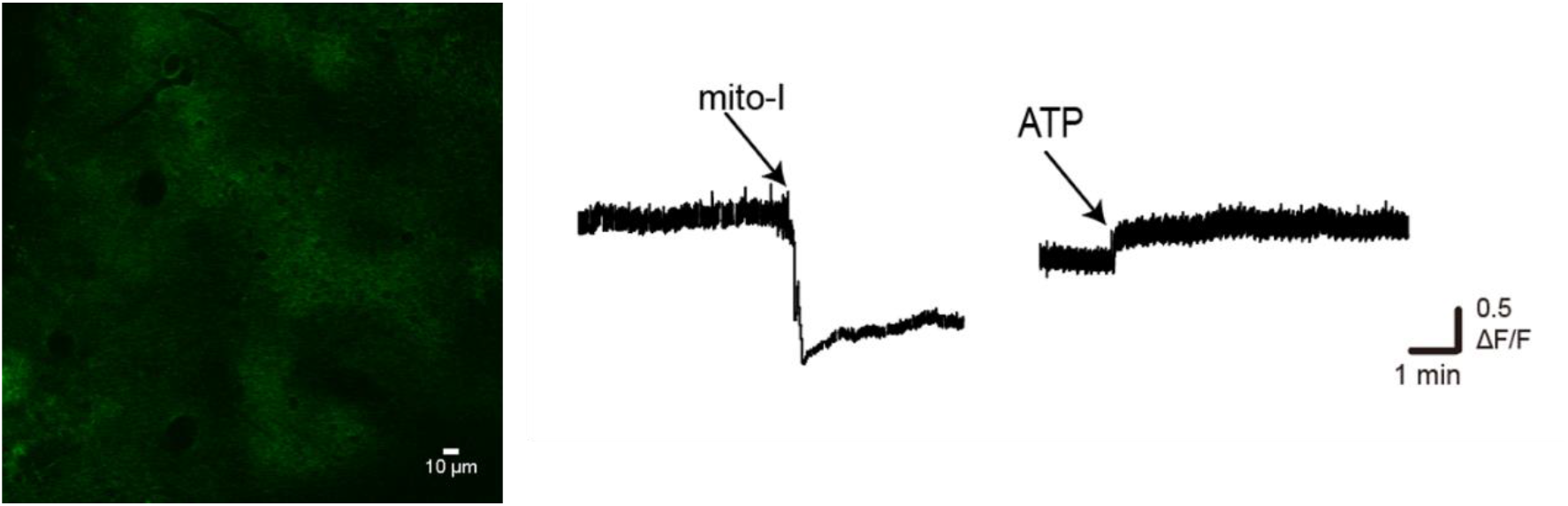
The fluorescence response to mito-I and ATP measured in neurons.

**Fig S1.**
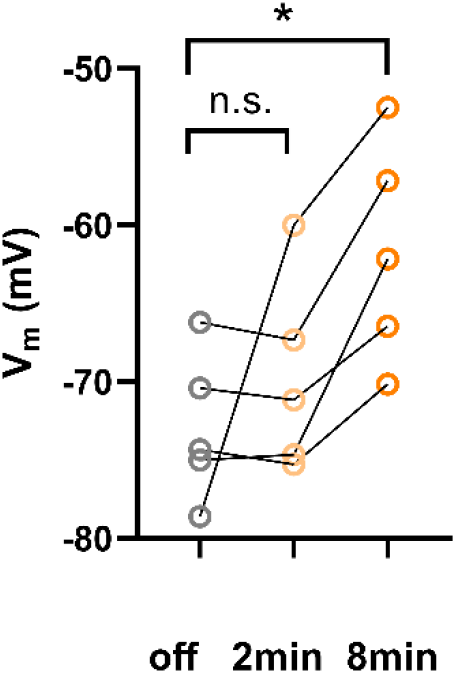
The changes in membrane potential caused by mito-I at different times.

**Fig S4.**
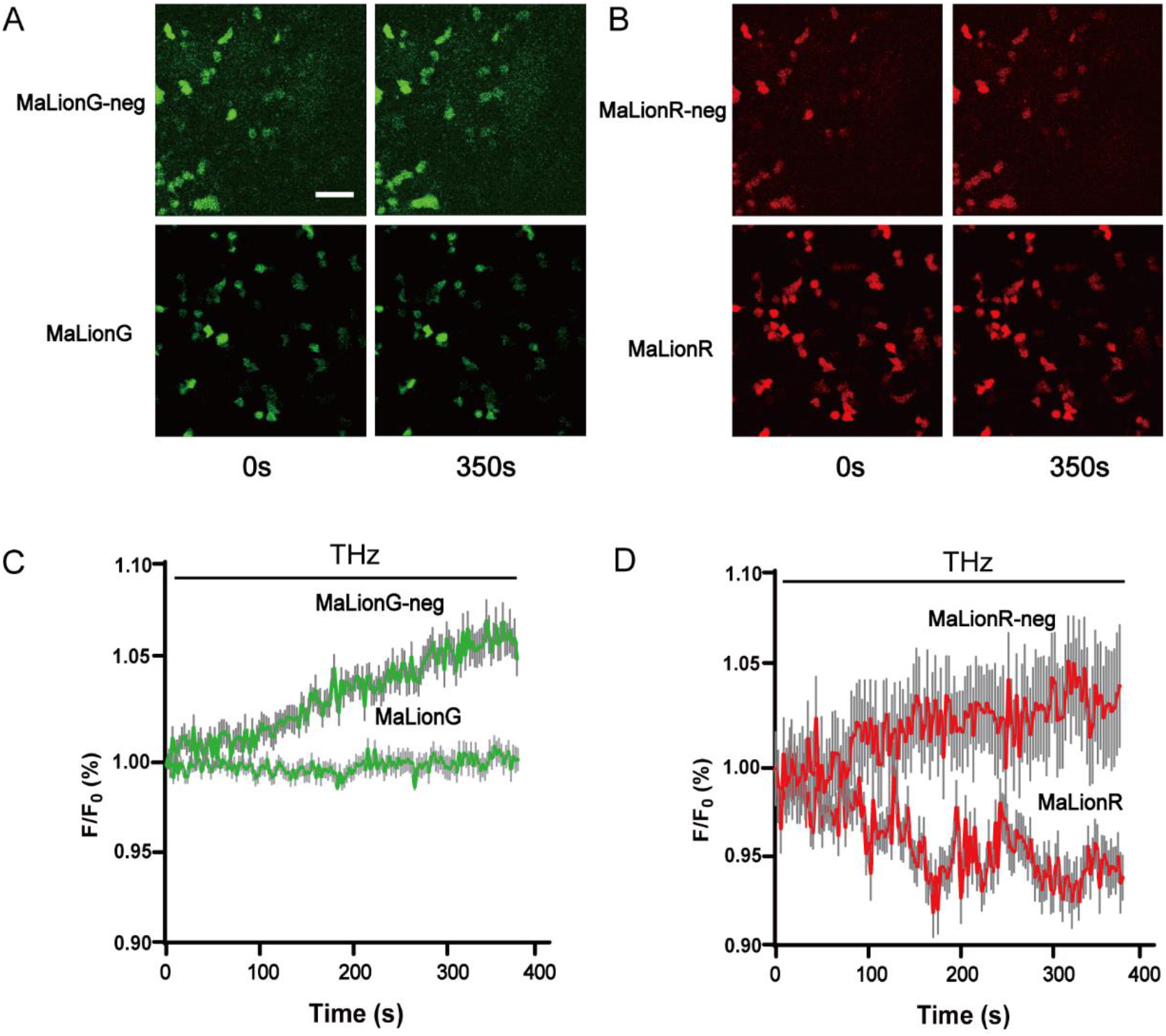
The changes of intracellular ATP during THz irradiation. **(A)** Fluorescence images of HEK293T expressing MaLionG and MaLionG-neg when the irradiation duration is 0s (left) and 350s (right) respectively. **(B)** Fluorescence images of HEK293T expressing MaLionR and MaLionR-neg when the irradiation duration is 0s (left) and 350s (right) respectively. **(C)** and **(D)** Normalized F/F0 was measured in MaLion and MaLion-neg expressing HEK293T cells in response to THz continuously applied (left: green; right: red); n=25 cells from 3 cultures. Scale bars represent 100 μm.

## Notes

### Competing Interest Statement

The authors have declared no competing interest.

### Summary of Updates

。

